# Mouse retinal cell behaviour in space and time using light sheet fluorescence microscopy

**DOI:** 10.1101/686626

**Authors:** Claudia Prahst, Parham Ashrafzadeh, Kyle Harrington, Lakshmi Venkatraman, Mark Richards, Ana Martins Russo, Kin-Sang Cho, Karen Chang, Thomas Mead, Dong Feng Chen, Douglas Richardson, Lena Claesson-Welsh, Claudio Franco, Katie Bentley

## Abstract

As the general population ages and the incidence of diabetes increases epidemically, more people are affected by eye diseases, such as retinopathies. It is therefore critical to improve imaging of eye disease mouse models. Here, we demonstrate that 1) rapid, quantitative 3D and 4D (time lapse) imaging of cellular and subcellular processes in the murine eye is feasible, with and without tissue clearing, using light-sheet fluorescent microscopy (LSFM) and 2) LSFM readily reveals new features of even well studied eye disease mouse models, such as the Oxygen-Induced Retinopathy (OIR) model. Through correlative LSFM-Confocal imaging we find that flat-mounting retinas for confocal microscopy significantly distorts tissue morphology. The minimized distortion with LSFM dramatically improved analysis of pathological vascular tufts in the OIR model revealing “knotted” morphologies, leading to a proposed new tuft nomenclature. Furthermore, live-imaging of OIR tuft formation revealed abnormal cell motility and altered filopodia dynamics. We conclude that quantitative 3D/4D LSFM imaging and analysis has the potential to advance our understanding of pathological processes in the eye, in particular neuro-vascular degenerative processes.

## Introduction

Eye diseases, such as diabetic retinopathy, age-related macular degeneration, cataract, and glaucoma are becoming increasingly common with the increased age of the general population. Although advances in understanding and treating eye diseases have been made, the molecular mechanisms involved are still not fully understood. We believe that is partially due to the inadequate ability to image eye tissue in its natural, spherical state, to reveal the many distinct layers with interacting cell types oriented differentially within or between the layers. Optical Coherence Tomography (OCT) is an established medical imaging technique that uses light to capture micrometer-resolution, three-dimensional images non-invasively, now widely used as a diagnostic tool (Srinivasan et al. 2006; Huber et al. 2009). Its main strength lies in revealing information on tissue depth preserving the eyes natural state. However, its limitation lies in not being able to provide a wide field of view, cellular or molecular information. Furthermore, being a non-fluorescent method, specific proteins cannot be labelled and tracked to investigate mechanism. Currently only confocal microscopy can deliver this detailed fluorescently labelled information (Del Toro et al. 2010), but the 3D nature of the tissue is likely distorted during flat-mounting and it is currently not known to what extent this might impact the obtained results. For instance, the vascular biology field is one clear example where these limitations can have a substantial impact. The mouse retina is a common model used to study vascular development and disease; confocal imaging approaches have been used to measure vessel morphology, vascular malformations, junctional organization, and pathological tuft formation (Gerhardt et al. 2003; Bentley et al. 2014; Stahl et al. 2010). Moreover, vessel diameters are now being used to predict blood flow (Bernabeu et al. 2014, Baeyens et al., 2016). Distortions arising from confocal flat-mounting could therefore have important ramifications for the overall conclusions of several studies.

Changes in cellular and tissue morphology are a hallmark of many eye diseases. For instance, retinopathy of prematurity and diabetic retinopathy are characterized by excessive, bulbous and leaky blood vessels that protrude out of their usual layered locations. These malformed vessels cause many problems including the generation of abnormal mechanical traction, which pulls on the different layers, eventually leading to detachment of the retina (Nentwich & Ulbig 2015; Hartnett 2015). Yet, very limited information has arisen on the conformation and morphogenesis mechanisms of these vascular tuft malformations, despite a wealth of confocal studies of the related Oxygen Induced Retinopathy (OIR) mouse model (Connor et al. 2009).

Another limitation of confocal microscopy for imaging of mouse retinal angiogenesis is the inability to perform live imaging of endothelial cell dynamics. Endothelial cells move and connect in highly dynamic, complex ways to generate the extensive vascular networks required to perfuse the retina over time (angiogenesis). Whilst live-imaging techniques of blood vessels exist, such as the dorsal skinfold chamber or the cranial window (Brown et al. 2010), it is not possible to translate those to study retinal angiogenesis, as the mouse pups are very small, making it hard to access the retina tissue. There are a small number of reports on *ex vivo* live-imaging of the retinal vasculature with confocal microscopy, but with limited success as flatmounting the retina for live-imaging takes time from dissection to culture, and it is necessary to physically force the tissue onto the membrane, disturbing local tissue arrangement and mechanics (Sawamiphak et al. 2010, Rezzola et al. 2013). Furthermore, photobleaching, phototoxicity and long acquisition times continue to remain an issue.

Recent advances in Light Sheet Fluorescence Microscopy (LSFM) have demonstrated its strength for allowing the rapid acquisition of optical sections through thick tissue samples such as mouse brains (Stelzer 2014). Instead of illuminating or scanning the whole sample through the imaging objective, as in wide-field or confocal microscopy, the sample is illuminated from the side with a thin sheet of light. Thus, in principle LSFM would require little interference with the original spherical eye tissue structure, avoiding distortion of the tissue with flat-mounting. Moreover, LSFM is becoming a gold-standard technique to perform live-imaging in whole organs/organisms because it permits imaging of thick tissue sections without disturbing the local environment, while also reducing photobleaching and phototoxicity (Stelzer 2014; Reynaud et al. 2014). Thus, here we investigate the feasibility, advantages and disadvantages of LSFM for imaging the mouse eye for development or disease studies. We present an optimized LSFM protocol to rapidly image neurovascular structures, across scales from the entire eye to subcellular components in mouse retinas. We investigate the pros and cons of LSFM imaging of vessels over standard confocal imaging techniques in early mouse pup retinas. Importantly, we also demonstrate the benefits of LSFM using the OIR mouse model, where we discover previously unappreciated new spatial arrangements of endothelial cells in the onset of vascular tuft malformations due to the improved undistorted, 3D and 4D imaging capabilities of LSFM.

We conclude that LSFM quantitative 3D/4D imaging and analysis has the potential to advance our understanding of healthy and pathological processes in the eye, with a particular relevance for the vascular and neurovascular biology fields, as well as Ophthalmology.

## Results

### LSFM enables rapid 3D imaging of mouse eyes, and in particular retinas, in their natural state

To visualize the retinal vasculature using epifluorescence or confocal microscopes, the retina is flat-mounted by making four incisions before adding a cover slip containing mounting medium onto glass slides (Fig. 1a, upper panel). To image samples using LSFM, however, samples are suspended in their natural state in low-melting agarose (Fig. 1a, lower panel). This enables imaging of the vasculature of large and intact samples such as the whole eyeball (minus the sclera and cornea) (Fig. 1b), the iris (Fig. 1c), or the optic nerve (Fig. 1d). Using LSFM, it was possible to observe the superficial, intermediate and deep vascular plexus (Fig. 1e and Suppl. Fig. 1b) of a retina in its native conformation (Fig. 1f). In contrast to imaging the retinal vasculature using the confocal microscope, where stacks contain 8 - 15 images, acquiring a stack of the retina using LSFM contains 200 - 300 images, but the imaging time is much shorter (∼60 sec vs. ∼10 min).

**Figure 1:**
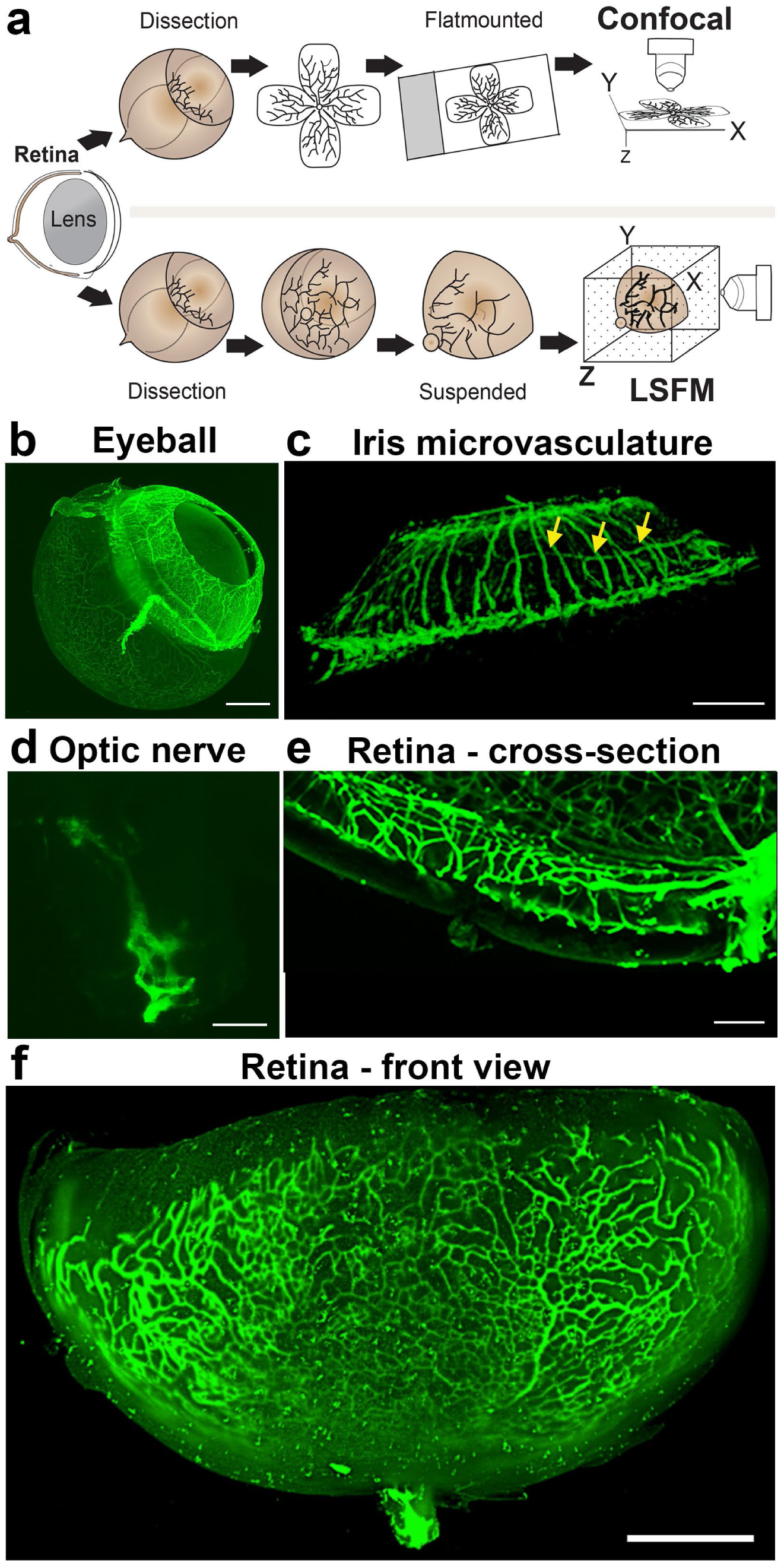
Imaging of the whole eye using light sheet microscopy. **a,** Schematic of retina preparation for imaging. For conventional confocal microscopy, four incisions are made to enable flat-mounting of the retina onto a cover slip. For LSFM, pieces of the retina are suspended and imaged from a right angle. **b,** Maximum intensity projection (MIP) of a P15 mouse eyeball (z = 274 slices). Vessels were visualized with IsoB4 staining. Scale bar, 500 µm. **c,** 3D-rendered image of the adult iris microvasculature (z = 263 slices). Vessels were visualized with IsoB4 (yellow arrows). Scale bar, 250 µm. **d,** MIP of the optic nerve. Vessels were visualized with IsoB4 staining (z = 176 slices). Scale bar, 50 µm. **e,** MIP of a cross section of a P10 mouse retina. Vessels were visualized with IsoB4 staining. Scale bar, 100 µm. **f,** MIP of a whole P10 mouse retina suspended and imaged intact (z = 176 slices). Vessels were visualized with IsoB4 staining. Scale bar, 500 µm. See supplementary figure 1 for additional images.

Imaging the iris microvasculature (Fig. 1c) surprisingly revealed that the vasculature network is immature at P15 (Fig. 1b, Suppl. Fig. 1), and that it remodels into a mature network in adulthood (Fig. 1c). A network of capillaries was visible at P15, whereas the adult microvasculature consisted of radial branches of small vessels and capillaries in a relatively linear pattern. The major arterial circles (MICs) around the iris root were developed in both P15 and adult mice (Fig. 1c and Suppl. Fig. 1a, yellow arrows). The images generated from adult mice using LSFM are consistent with a previous report using OCT to image the iris microvasculature (Choi et al. 2014). Using LSFM, the vessels appeared straighter, and the MICs were not as close to the iris root, which could be because OCT involves live-imaging of the vasculature, without mechanically removing the sclera and cornea.

We next tested whether LSFM could resolve subcellular structures in the retinal vasculature such as the Golgi apparatus, which has recently been shown to be important for inferring cell polarity during vessel regression (Franco, C.A. et al. 2015). We found it feasible to stain and image the Golgi organelle (Golph4, Alexa 647) and the collagen IV-containing basement membrane around the vessels (Suppl. Fig. 1c, d). On static images of mice expressing lifeAct-enhanced green fluorescent protein (EGFP) we performed deconvolution to reduce the light scattering effects and found this gave a marked improvement to the resolution of actin bundles within endothelial cells (Suppl. Fig. 1e-i). Taken together, LSFM can rapidly generate 3D images of the murine eye in its native form across scales, with tissue, cellular and subcellular resolution.

### LSFM enables concurrent imaging of retinal cell types within and between the retinal layers

A growing number of reports show that neurovascular interactions in the eye are important during development and disease progression (Akula et al. 2007; Narayanan et al., 2014; Nentwich and Ulbig, 2015; Usui et al. 2015; Verheyen et al. 2012). Neurons and vessels are however currently difficult to image in the same retinal tissue across all layers, without distortion due to the way they are arranged in the retina (Fig. 2a, b). Neurons are currently imaged by making vertical sections, orthogonal to the three vascular layers (the superficial, intermediate and deep plexus) (Fig. 2b “side view”), on paraffin or cryo-embedded retinas, which necessarily means losing the ability to observe vascular branching in the horizontal layer in the same tissue. Likewise, studies focused on the retinal vasculature use whole mount images of the retina viewed from above (Fig. 2b, “top view”) using horizontal optical sections (Usui et al. 2015), which does not allow proper imaging of retinal neurons spanning between the layers because of insufficient z-resolution in confocal microscopy. Thus, we next investigated whether concurrent imaging of neurons and vessels in the same sample might be achieved with the optical sectioning and rotational viewing capacity of LSFM.

**Figure 2:**
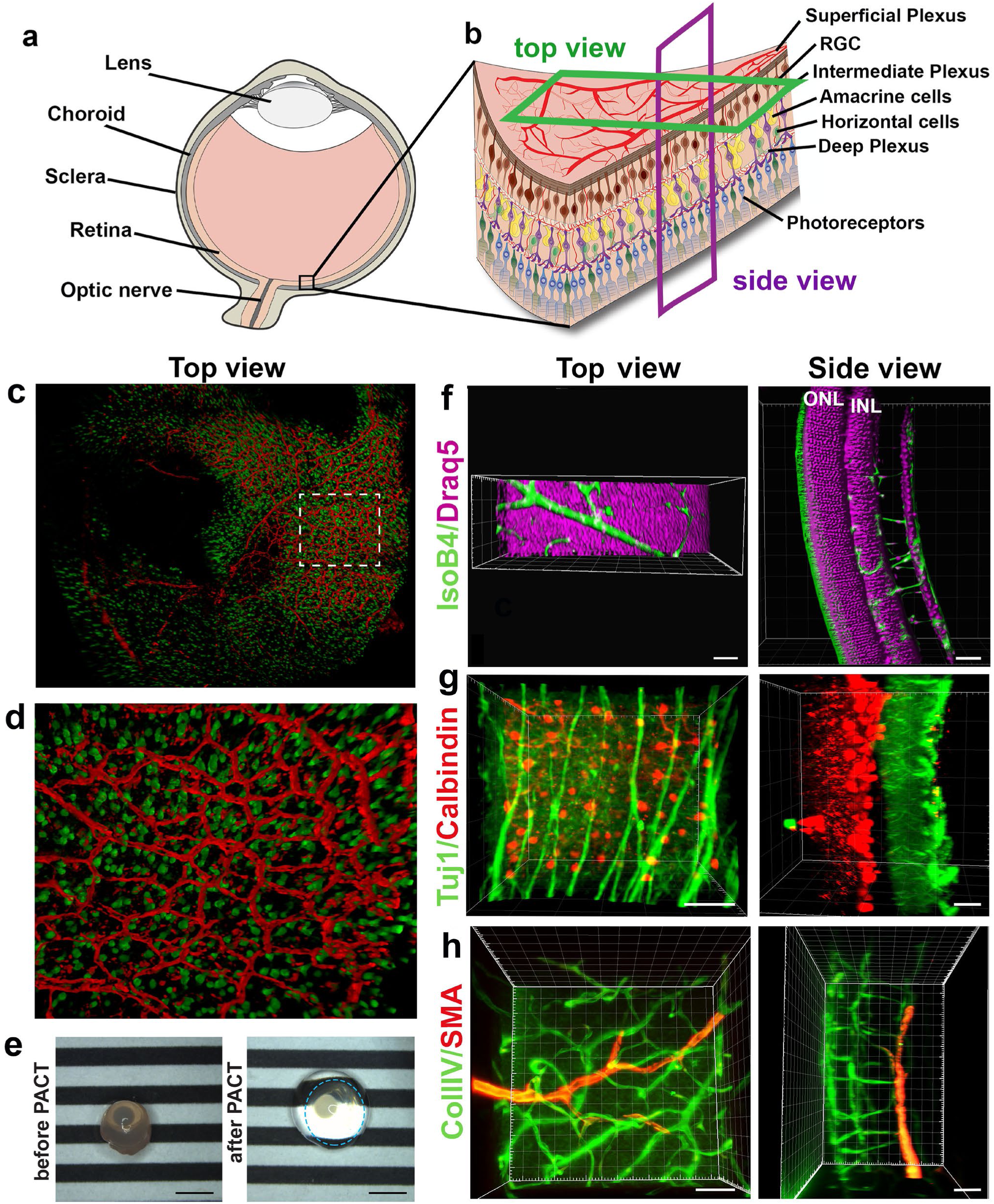
3D reconstruction of nerves and vessels in one image. **a,** Schematic of an eyeball. **b,** Schematic of the retina and its cell types. **c,** Retinal eye cups expressing yellow fluorescent protein, YFP (green) were harvest from Thy1-YFP mice and stained with Isolectin IB4 (red). The retinal eye cups were mounted and imaged with LSFM. **d,** enlarged region of c. **e,** Representative image of an eyeball before clearing (left panel), and an eyeball after PACT clearing (right panel). The circle around the cleared eyeball depicts the outline of the eyeball. Scale bar, 2 mm. **f,** Draq5 staining (magenta) visualizes the inner nuclear layer (INL) and outer nuclear layer (ONL) of the adult mouse PACT cleared retina. Vessels were visualized by IsoB4 staining (green). **g,** Tuj1 (green) and calbindin (red) visualize the ganglion and horizontal cells in the mouse PACT cleared retina. **h,** Smooth muscle actin (SMA, red) and Collagen IV staining (Coll.IV, green) visualize the three vascular layers and smooth muscle cells in the mouse PACT cleared retina. Scale bars, 50 µm.

We found that eye cups from P3 C57BL/6 Thy1-YFP mice, labelling retinal ganglion cells in yellow combined with IsolectinB4 labeled vasculature provided 3D high resolution images without the need for tissue clearing (Fig. 2c, d, Movie S1). However, we found that including lipid removal/permeabilization as part of a full tissue clearing protocol further improves resolution for eye cups at later stages of development, when more of the retinal layers have formed (Fig. 2e-h), as it decreases the scattered light caused by imaging thicker tissue with the light sheet (Richardson & Lichtman 2015).

We tested several different clearing methods to see which was better suited to retinal tissue. We first tested the aqueous-based clearing methods ScaleA2 and FRUIT (Hou et al. 2015; Hama et al. 2011). However, these clearing methods did not lead to higher quality images and made tissue-handling very difficult during imaging due to the high viscosity of the FRUIT clearing agent. We also tested the passive aqueous-based methods CUBIC-R (Kubota et al. 2017) and PROTOS (Murray et al. 2015), but again found little improvement. Since many studies use animals genetically engineered to express fluorescent markers such as Tomato or GFP, we decided not pursue solvent-based clearing methods such as iDISCO, which do not maintain fluorescent protein emission for more than a few days after the clearing process (Renier et al. 2014). Overall, we found PACT was the most efficient and effective clearing method for retinal tissue, likely because it is relatively thin (Yang et al. 2014; Treweek et al. 2015a). PACT cleared adult retinas with Draq5 staining, which stains all nuclei, visualizing the inner nuclear layer (INL) and outer nuclear layer (ONL) (Fig. 2f, Movie S2). The deep vascular plexus, visualized by IsolectinB4 staining could be seen between the ONL and INL, whereas the intermediate vascular plexus bordered the INL as expected. The superficial vascular plexus is located on the inner retinal surface together with nuclei of the retinal ganglion cells (Fig, 2f, Movie S2). Adult retinas were co-immunostained for Tuj1 and Calbindin, markers for retinal ganglion cells and horizontal cells, respectively. This immunostaining made it possible to appreciate the distance between these two cell types in the fully developed retina (Fig. 2g, Movie S3). 3D-rendered images of co-staining for smooth muscle actin and collagenIV moreover showed arteries of the superficial vascular plexus covered with smooth muscle cells (Fig. 2h, Movie S4).

### Vessel distortion due to confocal flatmounting revealed by correlative LSFM-Confocal imaging

As vascular measurements taken from confocal images are used as the standard for inferring the actual sizes of vascular structures in the retina, we next aimed to systematically quantify the 3D distortion of vascular structures incurred by flat-mounting and confocal imaging. In order to make direct, quantitative comparisons of the relatively small vessels in the superficial plexus, we used a correlative LSFM-Confocal approach: we first imaged the retinal tissue with LSFM, which retains the natural tissue curvature, then we melted the gel and flat-mounted the same retina onto a coverslip and imaged it again using confocal microscopy (Fig 3a). We first analyzed the largest vessels near the optic nerve and then smaller capillaries in the sprouting vascular front from P4 wt retinas. Images obtained with our correlative LSFM-confocal approach were then brightness/contrast adjusted and cropped and surface rendered using Imaris to focus on small regions of same vessel segments in the corresponding confocal and LSFM images. Dramatically shallower side views and cross-sections of vessels were evident in the confocal images compared to LSFM (Fig. 3b). We next quantified this shift in aspect ratio by measuring the diameter taken across the vessel in XY (hereafter “Width”) and down through the Z-axis (hereafter “Depth”) in confocal (Fig. 3c). For LSFM images, given the tissue can be at any orientation in the agarose with respect to the objective, the XYZ coordinate system of the image stack is not indicative of the equivalent width/depth measurement in confocal. Therefore, the orientation of the surrounding vascular plexus at the point of the vessel segment was used as a reference surface “plexus plane” to make the corresponding “width” diameter measurement, as it is equivalent to the XY plane in the corresponding confocal image. Similarly, the “depth” diameter in LSFM was defined as perpendicular to the plexus plane and width measurement (equivalent to the diameter through the z-stack in confocal). Vessels were significantly more elliptical (wider and shallower) under confocal than LSFM indicative of being compressed during flat-mounting (Fig. 3d,e).

**Figure 3.**
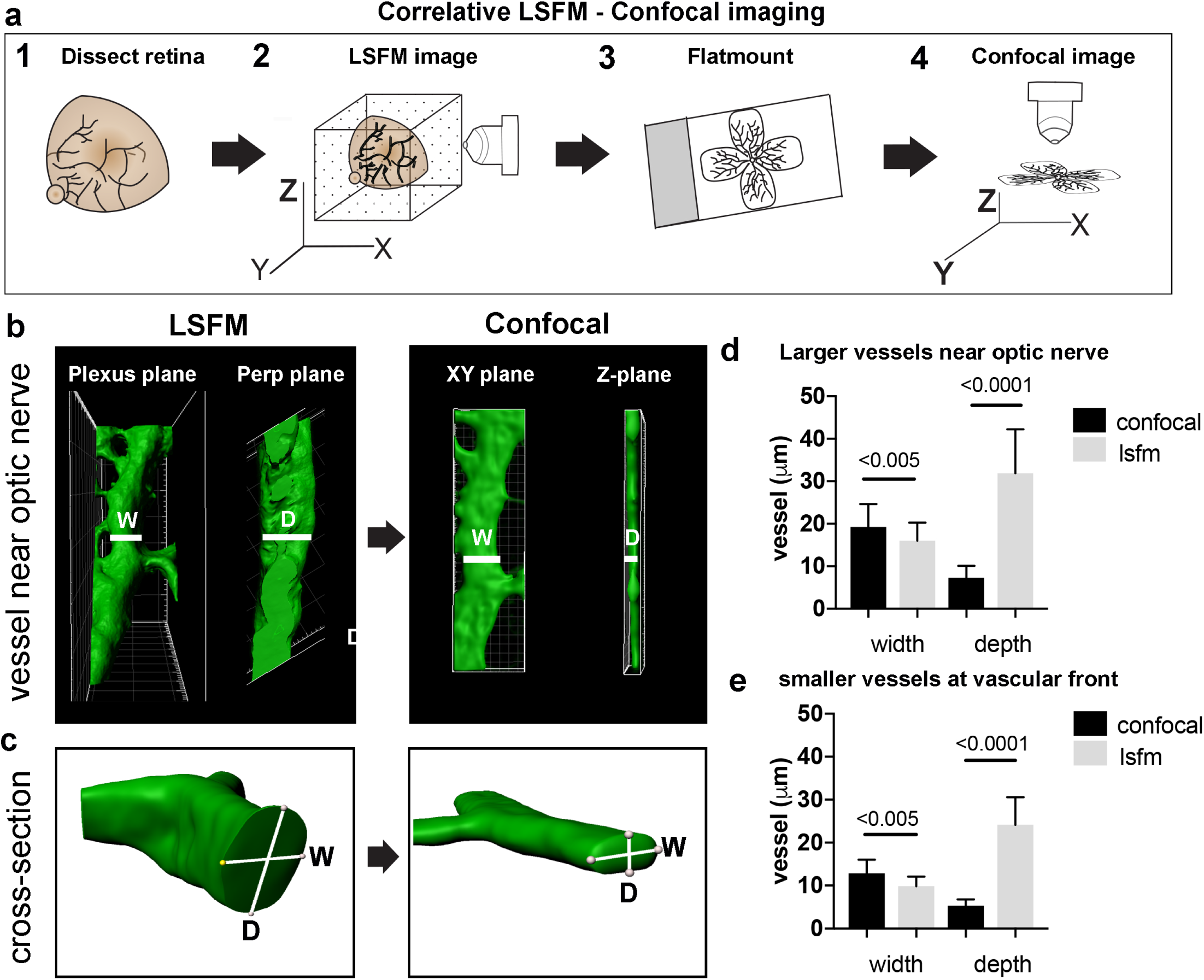
Vessel depth distortion in confocal due to flatmounting. **a,** Schematic showing the correlative LSFM-Confocal imaging approach used to quantify vessel distortion incurred by flatmounting. **b,** the same large vessel segment imaged first with LSFM then confocal (surface rendered in Imaris). By orienting with the surrounding vessel connections to determine the plexus plane (equivalent to the XY plane in confocal) and the plane perpendicular to it (“perp plane”), which is equivalent to the Z plane in confocal, comparative width (W) and depth (D) measurements can be made of the same vessel segment. **c,** cross sectional views of another representative large vessel near the optic nerve shows how the aspect ratio of W and D is shifted to an ellipse in confocal. Near Optic: n = 60 vessels from 6 retinas (7 images). Vascular Front n = 28 vessels in from 4 retinas (4 images) for each confocal and lsfm.

### LSFM enables 4D live-imaging with subcellular resolution, revealing rapid, transient “kiss and run” tip cell adhesions at the sprouting front

*Ex vivo* live-imaging could be a useful tool to study tip cell guidance during the angiogenic sprouting process in the mouse retina, but it has proven to be challenging with conventional microscopy. Existing *ex vivo* confocal methods to live-image retinal vasculature, tissue handling leads to damage of the tissue, as it involves either flat-mounting the retinas onto a membrane and then submerging it in medium (Sawamiphak et al. 2010), or cutting the retina into fragments and embedding them in fibrin gels prior to imaging (Rezzola et al. 2013).

We therefore established a protocol for live-imaging of the growing retinal vasculature using LSFM. We first crossed mT/mG mice with Cdh5(PAC)-CreERT2 mice and injected them with tamoxifen to induce endothelial GFP expression (Muzumdar et al. 2007; 2010; Wang et al. 2010). Surprisingly, connections between ECs formed very rapidly (within 20 min) and regressed just as rapidly (Fig. 4a, Movie S5). Such transient “kiss and run” adhesion and release style interactions between ECs (as opposed to full adhesions or anastomoses, where the connections stably remain) have only been previously reported in glycolysis-deficient ECs *in vitro* with “normal” connections being more stable *in vitro* (Schoors et al. 2014). The dynamics *in vivo* were therefore assumed to be slower and more stable than *in vitro* live-imaging, however our new observations indicate a very different set of dynamics and inter-cellular behaviors may be at work in the complex *in vivo* tissue. Timing is crucial, as the VEGF gradient dissipates after the retinas are dissected and submerged in agarose, at room air and the tissue is therefore no longer hypoxic. However, the directed growth of the filopodia towards the vascular front in our LSFM movies suggests that this gradient remains intact for at least the first few hours after dissection. We found filopodia length and extension, retraction dynamics quantification feasible (Fig. 7d,e,f). Further back from the sprouting front, in the vascular plexus (Fig. 4b, Movie S6), we occasionally observed the formation of connections over the course of a few hours, however, branch formation was a rare occurrence. Notably, we did not observe EC apoptosis under these imaging conditions indicating conditions are viable.

**Figure 4:**
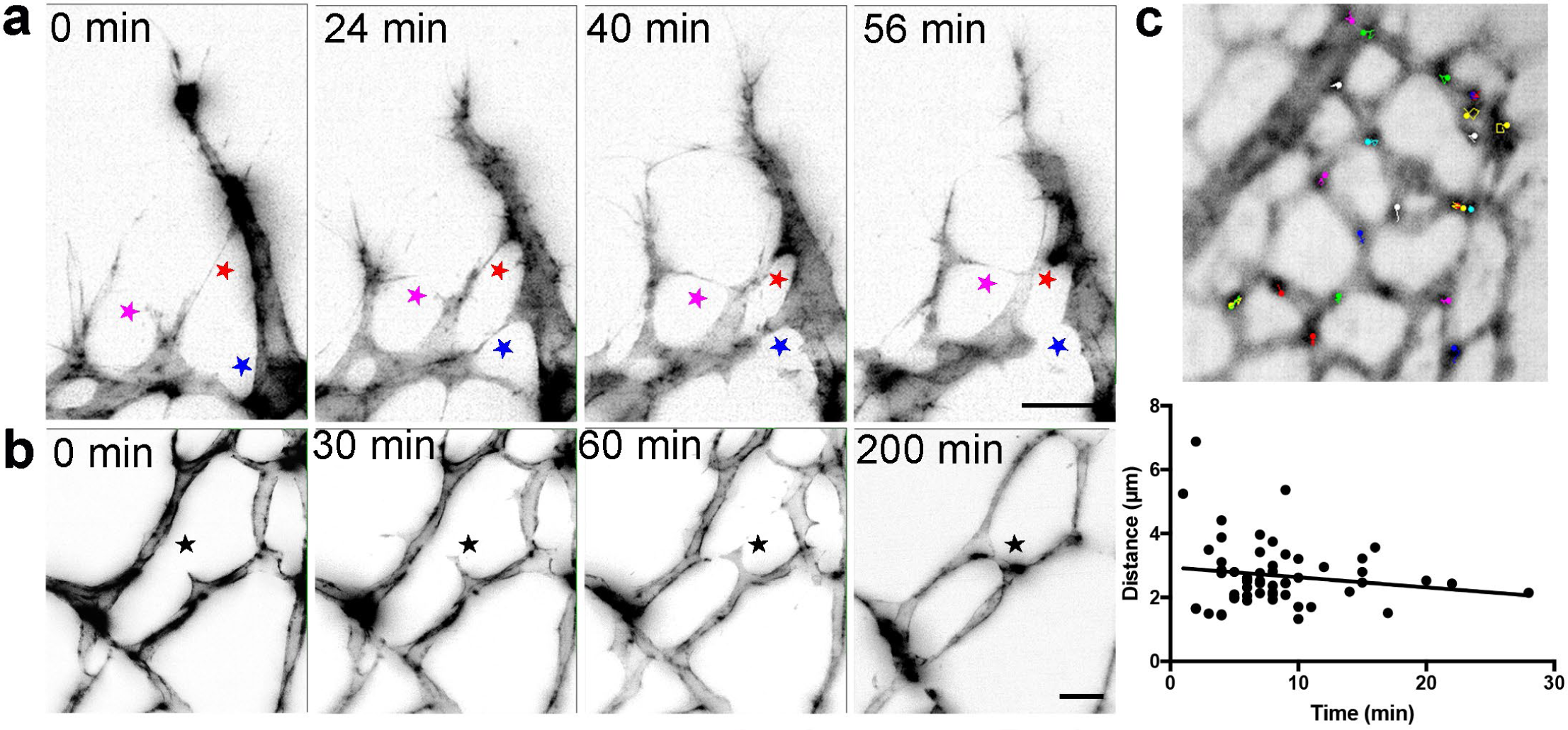
Live-imaging of the retinal vasculature. **a,** Single maximum intensity projections (MIP) of an hour time lapse movie show long, slender filopodia, and rapid fusion and disconnection of tip cells at the vascular front of mT/mG x cdh5 (PAC) CreERT2 mice (stars). **b,** MIPs of a time lapse movie reveal the connection between two branches in the capillary plexus (star).

We next assessed the feasibility of using LSFM to live-image intracellular processes. We dynamically imaged mice expressing lifeAct-EGFP, which fluorescently labels F-actin (Fraccaroli et al. 2012) with LSFM and quantified the movements of actin-enriched bundles within the endothelial cell bodies in the sprouting front during developmental angiogenesis. Quantitative subcellular actin live-imaging was found feasible (n=6 retinas) with the average distance travelled by each bundle found to be 2.56 µm (Fig. 4c, movies S7, S8 and S9).

Taken together, our LSFM permits the visualization in real-time of cellular movements with subcellular resolution in the mouse retina, with minimal distortion.

### Three subclasses of pathological retinal neovascular tufts revealed with LSFM

We next sought to image vessels that have grown pathologically, in order to determine whether this imaging method could be used to gain greater insights into eye disease. To this end, we used the OIR model, where mouse pups are placed in 75% oxygen from P7 to P12, and are then kept at room air from P12 to P17 (Connor et al. 2009). During the hyperoxia phase, the vasculature regresses, and in the subsequent normoxia phase, new vessels grow in an abnormally enlarged and tortuous manner (Connor et al. 2009). Furthermore, vessels also start to grow into the vitreal space forming bulbous vessels, known as “vascular tufts”, above the superficial vascular layer (Fig. 5a). In the past, it has been difficult to analyze and characterize the growth of these tufts because they are large formations which appear to be distorted by the flat-mounting process. By performing IsolectinB4 and ERG immunostaining to visualize endothelial cells (ECs) and their nuclei, we obtained 3D-reconstructions of the tufts and were able to first classify them into different groups by quantifying both volume and number of nuclei (Fig. 5a,e). As expected, we found that the number of nuclei increased with the size of the tuft (R^2^ = 0.83). Interestingly, however we found many small tufts, and only very few large tufts. The smallest tuft we could identify had two nuclei parallel to each other, the cells growing straight up into the vitreous (Fig. 5a, upper panel, Movie S10). We found that most tufts have between 4 and 20 nuclei (“Medium tufts”, Fig. 5a, second panel row, Movie S11). We also identified a few very “large tufts” with over 20 nuclei (Fig. 5a, third panel row, Movie S12). Next, we quantified the number of connections between the vasculature and the tuft (Fig. 5b). The large tufts had a higher number of connections to the existing vasculature (R^2^ = 0.61), however surprisingly for medium and large tufts, despite very different nuclei counts, the tuft volume and number of connections to the plexus remain approximately constant (Fig. 5c). However, the number of connections and tuft volume transition sharply to ∼2.5 fold and ∼3 fold respectively, when the number of nuclei in the tuft exceeds twenty. This indicates that proliferation or an influx of cells to the tuft does not increase tuft volume, but rather tuft volume only significantly increases when the number of connections to the plexus also increases. Based on this observation, we propose that large tufts are in fact formed by fusion of 2 or 3 medium tufts.

**Figure 5:**
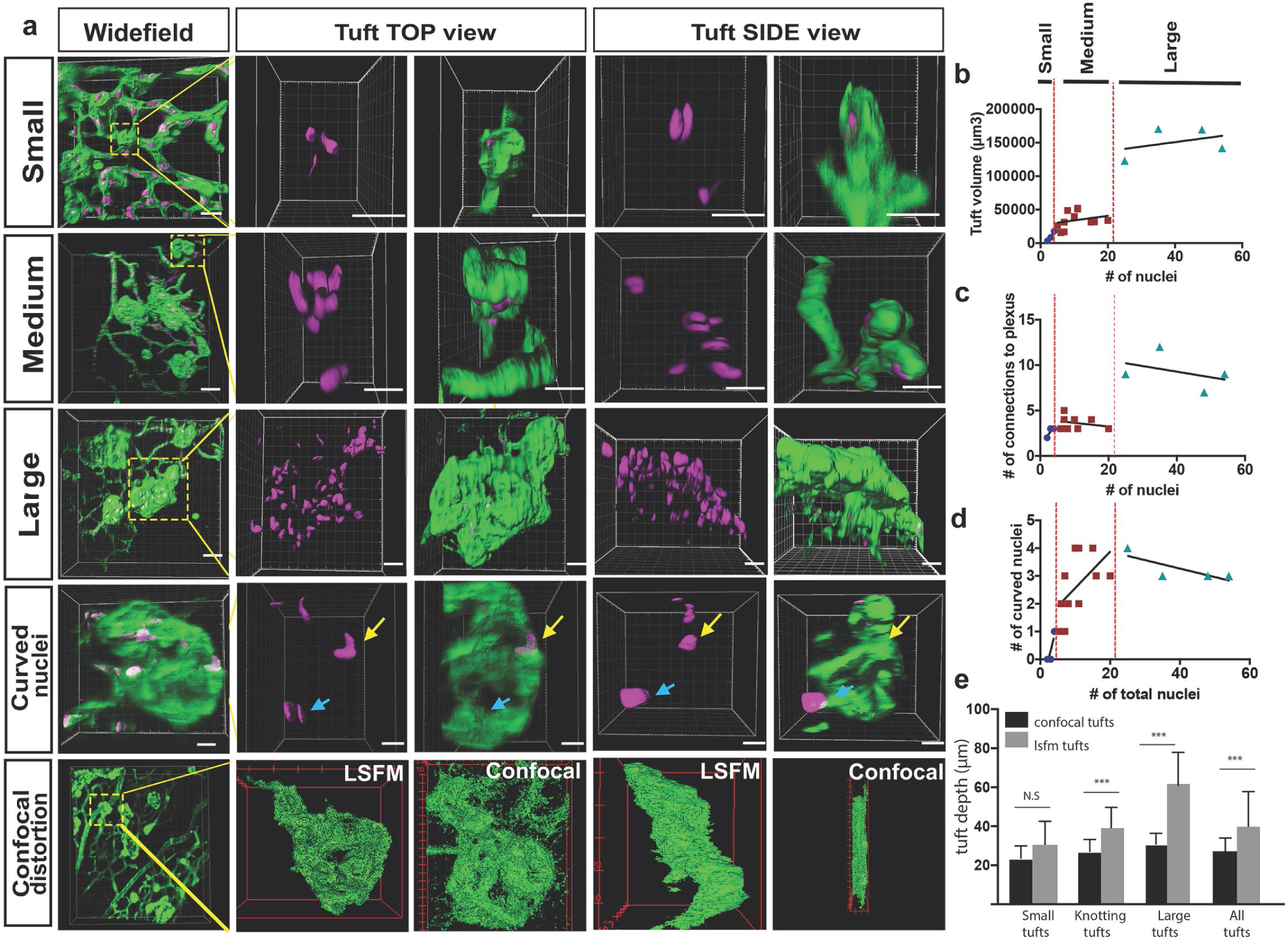
Analysis of three subclasses of OIR vascular tuft. **a,** Representative 3D-rendered LSFM images of small (1^st^ row), medium (2^nd^ row), and large (3^rd^ row) tufts showing the vasculature (IsoB4, green) and endothelial nuclei (ERG, magenta), scale bar = 10 µm. For all widefield images, scale bar = 40 µm, yellow box indicates tuft in situ; 4^th^ row **-** representative 3D-rendered images showing curved nuclei in a medium tuft, yellow arrows indicate curved nuclei, blue arrows indicate flat nuclei parallel to each other. Scale bar, 10 um. 5^th^ row: correlative LSFM-confocal microscopy of the same tuft reveals the tuft depth distortion (side view) incurred with confocal flatmounting versus LSFM. **b,** The volume of the tufts versus the number of nuclei per tuft. **c,** The number of vessel connections between the tuft and the underlying vascular plexus versus the number of nuclei. **d,** Quantification of the number of curved nuclei per tuft versus the total number of total nuclei per tuft. **e,** quantifications of tuft depths per subclass size in LSFM vs confocal images, significant difference shown using unpaired t test, *** means P <0.0001.

We observed that some of the vascular tufts contained highly curved nuclei (Fig. 5a, fourth panel row, yellow arrow, Movie S13). Quantification of the number of curved nuclei/total nuclei in a tuft showed that in small and medium tufts, the number of curved nuclei correlated well with the number of total nuclei (Fig. 5d). In large tufts (over 20 total nuclei), the number of curved nuclei was stable suggesting actually a decline in curved nuclei as the number of cells in the tuft increased. Thus, the relative number of curved nuclei per tuft could also be used as a clear marker to distinguish medium and large tufts. As curved nuclei indicate cells are under severe mechanical strain, twisting or turning them around (Yuntao et al. 2018), this suggests that larger tufts may be more stable and mature, whereas the small and medium ones are under more tension, still forming with significant forces curving and pulling the cells around in the tuft. Interestingly, highly curved nuclei have been shown to result in rupture of the nucleus and DNA damage (Yuntao et al. 2018), which may further exacerbate dysfunctional cell behavior in tuft formation. It should be noted that care should be taken to rotate the image stack to confirm nuclear curvature, as two nuclei parallel to each other can look like only one nucleus (Fig. 5a, fourth panel row, blue arrow), emphasizing the importance of 3D imaging with LSFM as rotating and viewing tufts from the side without distortion is not possible with confocal.

Finally, to quantify the level of distortion of vascular tufts incurred by flat-mounting and confocal imaging, we compared tuft depth measurements between retinas imaged with confocal and LSFM (depth defined the tuft length orientated perpendicular to plexus plane). The change in depth was particularly striking and more pronounced for larger tuft structures (Fig 5a bottom panels, e), further confirming that LSFM is superior to confocal to image larger structures in the eye.

### LSFM OIR Case-study: Pathological retinal neovascular tufts have a knotted morphology

In order to gain better resolution to characterize the specific morphology of the different sized tufts we next performed computational image deconvolution on cropped LSFM images of vascular tufts (see Materials and Methods), which helped to decrease the scattered light caused by imaging thicker tissue with the light sheet without the need to clear the tissue (Richardson & Lichtman 2015). Upon deconvolution a dramatic and previously unappreciated “knotted” morphology of the tufts was evident across all tuft classes; often tufts had one or more holes going through (Fig. 6a,b white arrows. Also see Suppl. Fig. 2a,b for more rotational views and original rotational movies S14,S15). To describe these 3D tuft structures in detail, we next explored three systematic approaches: 1) by slowly shifting clipping planes through the tuft from the vitreous facing side to the plexus-connecting side of the tuft it was possible to better appreciate the upper and lower 3D organization of the tuft; 2) carefully rotating and hand-drawing the tufts surface rendered structures from every angle and 3) comparing the colour-labeled positions of nuclei to indicate their depth position in the tuft. The first approach revealed that the tuft shown in Fig. 5a fourth panel row, had a figure of eight knot, with two clear holes through the tuft and an unexpected vessel connecting the upper vitreous surface of the tuft to the plexus (Fig. 6c and Suppl. Fig. 2b, blue stars, Movie S13). A combination of the second two approaches revealed a swirl structure to two tufts (small and medium in size) akin to a snake coiling upon itself, with several highly curved nuclei (Fig. 6d-I, Movies S16, S17) and a central hole through the entire tuft. The smaller swirled tuft had two layers, while the medium tuft had three. In both cases the top, vitreous facing surface of the tuft appeared sprout-like with protrusive shapes, a morphology consistent across many tufts, e.g. Suppl. Fig. 2c white arrow and Fig. 7a. Overall, all three approaches were extremely useful for better interpreting these complex 3D structures, providing a much deeper understanding of them than viewing as maximum 2D projections or simply rotated on a screen.

**Figure 6.**
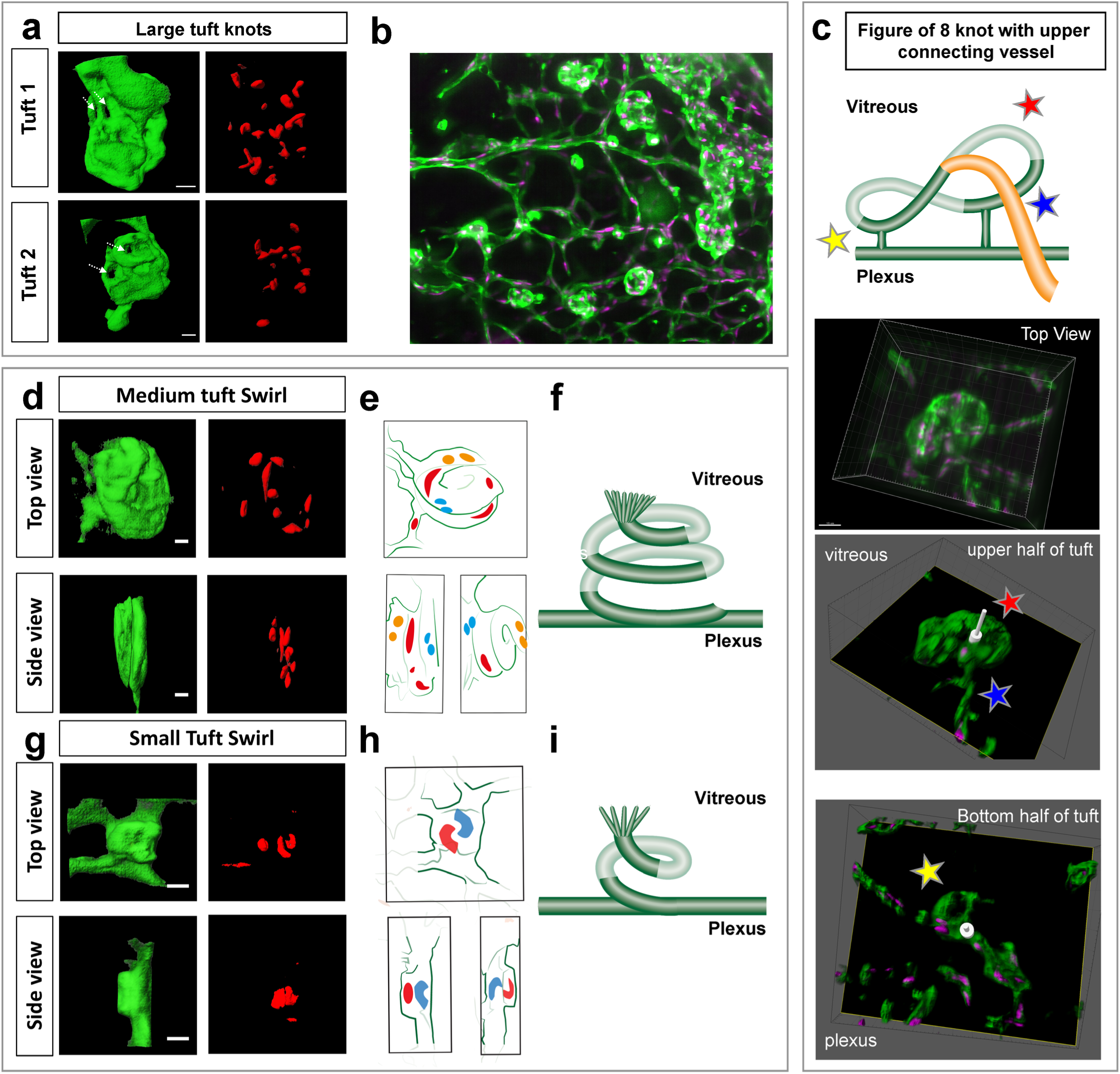
Knotted morphology of neovascular tufts revealed with LSFM. **a,d,g,** representative 3D-rendered images (generated using IMARIS software) of large (upper panel), medium and small tufts showing the vasculature (IsoB4, green) and endothelial nuclei (ERG, red) from rotational movies S14,15 See Supp. Fig. 2a for further views from different angles of the large tufts. White dashed arrows indicates a hole through the tuft. Scale bar, 30 µm. **b,** widefield LSFM of OIR retina demonstrates that the knotted morphology is hard to discern from afar. **c,** detailed 3D clipping plane and 3D rotational drawings of an individual knot reveal a figure of eight structure with two clear holes through the tuft aswell a vessel connecting from the upper, vitreous facing surface of the tuft to the plexus below (blue star). Stars mark corresponding regions from the illustration to the images - lower tuft loop nearer plexus (yellow star), upper tuft loop nearer vitreous (red star). See also Suppl. Fig. 2b for detailed 3D drawings made from each rotational view of this tuft with clipping planes, and Movie S13. **e,h,** 3D sketches made from rotational movies S16,17 to better elucidate nuclei: blue nuclei - bottom of tuft (near plexus), red nuclei – middle of tuft (in **e**), top of tuft (facing vitreous) in **h,** yellow nuclei - top of tuft (facing vitreous) in **e,f,i,** schematic illustrating the swirling tuft morphology observed in **d-h** with three layers for the medium tuft (**f**) and two for the small one (**i**).

### 4D LSFM live-imaging of the OIR mouse model reveals altered cell dynamics

To gain insights into endothelial cell behavior in vascular tufts, we next imaged the OIR-induced tufts dynamically with LSFM. Thereby, we observed that filopodia extended/retracted from abnormal vascular tufts, similar to what is seen in the extending vascular front during development of the retina vasculature. However, filopodia formed from vascular tufts remained very short (mean 4.3 µm) as compared to normoxia (mean 14.84 µm) (Fig. 7a, d). In OIR, filopodia more rapidly extended and retracted, without making connections (Fig. 7a, e, f, Movie S18). As the VEGF gradient is expected to be disrupted in the OIR model, timing from dissection to imaging is not as crucial. However, most filopodia movements occurred in the first few hours under this pathological condition. When imaging other parts of the OIR retinas to the tufts, we observed intriguing, abnormal EC behaviour. Their movements were undirected and appeared to involve blebbing-based motility (Fig. 7b, c, Movie S20). We observed both cells that were dividing, and undergoing apoptosis (Fig. 7b, Movie S19), which was not observed during normal conditions. This first live imaging of altered cell behavior in the OIR mouse model further highlights the potential of LSFM for new insights into disease processes.

**Figure 7.**
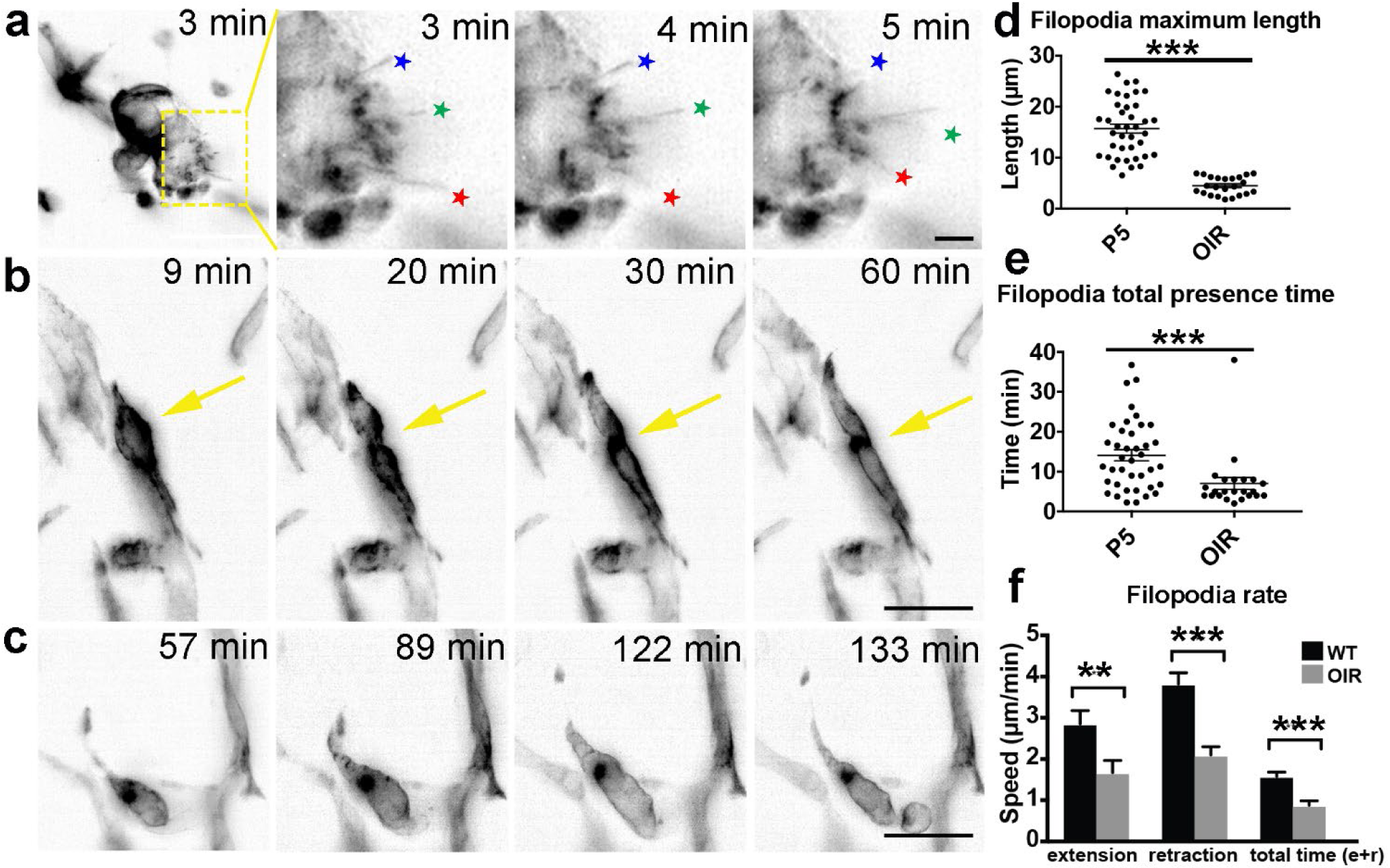
OIR live imaging **a,** Maximum intensity projections (MIPs) of a time lapse movie of a retinal tuft of a mouse in the oxygen-induced retinopathy (OIR) model visualize short, rapidly extending and retracting filopodia as compared to control retinas (stars). **b,** MIPs of a time lapse movie of mouse retinal vasculature in the OIR model reveal cell shuffling in real-time (arrow). **c,** MIPs of a time lapse movie of mouse retinal vasculature in the OIR model reveal abnormal vessel growth in real-time. Scale bar, 20 µm. **d,** the maximum length that each filopodia reached was measured for each filopodia over time in P5 and OIR conditions. **e,** the total time that each filopodia was present during the experiment; this time is calculated from when one filopodia appeared and then disappeared. **f,** speed of extension and retraction of filopodia were calculated for P5 and OIR conditions. Total n= 67 and 23 filopodia in 8 and 3 cropped movies from 3 independent P5 and 1 OIR experiment.

## Conclusions and Discussion

Although LSFM is becoming increasingly popular, studies to date have not attempted to use it to image eye tissue. We have therefore established the first protocols to image and clear mouse eye tissue using LSFM. Because this protocol utilizes optical sectioning of whole mount tissue, we found that LSFM is a very useful tool to rapidly image and reveal eye tissue at cellular and subcellular resolution without distortion of the sample due to flat-mounting, with the benefit to view and rotate structures in full 3D. As such, the present study provides a highly relevant and improved approach to examine the inter-relationships of normal neurovascular structures and the complex morphology of aberrant vascular structures in disease models, revealing for the first time a knotted morphology to the vascular tufts in OIR. We have also established a clearing protocol for eye tissue. We have furthermore established an *ex vivo* 4D live-imaging method to follow angiogenic growth in the mouse retina in real-time, both during development and under pathological conditions, and quantified that these dynamics are significantly altered in pathological conditions. The acquisition of 3D images of vascular structures at high spatial and temporal resolution within intact ocular tissue is both novel and significant. Overall, we strongly suggest the use of LSFM for 1) the study of larger or more complex 3D tissue structures reaching across the typical retinal layers, which are liable to distortion with standard approaches and 2) dynamic cell and subcellular processes in the mouse eye. We see far-reaching potential of the approach for deeper insights into eye disease mouse models in particular. For example, it would now be feasible to skeletonize larger portions of the vascular network (ultimately, the entire retina vasculature) in order to perform flow simulations and understand how the biomechanical feedback of flow impacts vessel growth in healthy and diseased eyes.

### LSFM vs Confocal: A Balanced Discussion

#### Benefits of LSFM

In general image acquisition with LSFM is widely known to be far faster than confocal due to the illumination of the entire optical plane at once combined with the use of a camera instead of detectors, as noted earlier, an extensive stack of the entire mouse retina can be acquired very quickly using LSFM (∼60 sec). 1) LSFM is better for imaging thicker or very large tissues (such as the eye cup, which is thin, but topologically spherical), due to the fast acquisition rates and the large, rotatable sample holder, removing the limited single view point from above with upright microscopes and slide mounting, 2) the illuminated plane generates less photobleaching and faster time frame rates for high temporal resolution live imaging of 3D / very thick tissues. We find LSFM imaging of the retina to be particularly informative over standard confocal microscopy when studying the following specific complex 3D and/or dynamic structures in the eye: 1<u>) the adult retina in full</u> - it is possible to visualize all three vascular layers in the LSFM, including direct cross-sectional viewing of the diving vessels oriented between layers by rotating the sample relative to the objective, which is not possible with confocal. Similarly, the iris and optic nerve can be observed in full, from any angle, undistorted with LSFM. 2) <u>abnormal enlarged vessels/tufts</u> - a newly characterized knotted morphological structure of tufts was revealed due to the improved 3D imaging and rotational viewing possible with LSFM. With confocal imaging the tuft shape can only be inferred from above and we found the depths were significantly distorted and compressed, which is likely why knots have not been described before. Interestingly the VE-cadherin staining of endothelial junctions of several OIR tufts shown in (Bentley et al 2014) indicated there were “holes” through tufts, as no junctional stains were found in clear pillars through them. However, the holes were not easy to confirm by isolectinB4 staining in the same samples, likely due to spreading of the vascular structure when it was distorted during flat-mounting. We can confirm here with LSFM there are holes through tufts and that tufts appear to consist of one or more long vessel structures intertwined, swirled and potentially looped upon themselves. 3) <u>Neurovascular interactions in one sample</u>, as neurons and vessels are oriented perpendicular to each other through the retina, they are normally imaged with separate physical sectioning or flattening techniques in either direction, prohibiting their concurrent observation. Optical sectioning of thick tissue and then rotating the undistorted image stacks allows both to be imaged for the first time together. Indeed, obtaining such images from one sample with LSFM will permit the quantification of vessels protruding through the neuronal layers, which is now only possible by performing time-consuming serial block-face scanning electron microscopy (Denk & Horstmann 2004). 4<u>) Live-imaging of developing mouse retinas</u> – this has proven very difficult with *in vitro* methods providing more reliable assays, e.g. embryoid bodies (Kearney & Bautch 2003; Jakobsson et al. 2010). Although embryoid bodies contain some other cell types such as pericytes, they still do not fully reflect the complex and tissue specific *in vivo* retinal environment. Importantly, the vessel-like structure formed in the embryoid bodies have never experienced flow. Moreover, the embryoid bodies are treated with VEGF supplied to the culture medium, while in vivo, endothelial cells are exposed to a VEGF gradient from the astrocyte network below. Our images suggest that the VEGF gradient remains intact in the retina samples for several hours in LSFM imaging. Furthermore, our protocol enables us to follow and quantify filopodia movements from minute to minute, revealing movements never seen before. Thus, we observed astonishing abnormal cellular and subcellular level dynamics under pathological OIR conditions by 4D live LSFM imaging.

#### Benefits of confocal over LSFM

Confocal microscopy has a fundamentally higher spatial resolution with less light scatter than LSFM; clearer, more precise images of smaller structures can be obtained, such as endothelial junctions and tip cell filopodia morphology provided the tissue sample is amenable to flat mounting without distortion or loss of information – i.e. it is naturally thin cross-sectionally and structures of interest have their main features in the XY plane, not in Z, XZ or YZ. Thus, confocal static imaging of normal developing vessels in a single layer of the retina will still yield better resolution images than LSFM and is very reliable for XY based quantifications such as branch point analysis. However, we find it is not reliable for acquiring accurate quantifications involving depth through Z such as vessel diameters or the morphology of cells that span between the layers (e.g. in the XZ or YZ planes). Thus, overall LSFM is not suggested to replace confocal for static developmental angiogenesis studies. However, to study and measure precise morphological attributes or dynamics of vessels with inherently 3D nature such as vessel radii, enlargements, malformations, diving vessels, iris, optic nerve or the deeper layers we find strong evidence to favour LSFM over confocal imaging. In general, the quantification time was comparable between LSFM and confocal images but there is potential for image analysis to require more effort for LSFM as files can become quickly large due to the rapid imaging (∼200GB for static imaging and up to 4TB for live multichannel imaging). If an older eye is being imaged the three vascular layers will be somewhat visually overlapping (e.g. in Supp. Fig. 1b), which could be hard to manually untangle due to the curvature, and as such represents a limitation. The preservation of the tissue depth information in the large z stack however, means by computationally fitting to the local curvature of the eye tissue one could computationally colour code and subtract the retinal layers out for independent viewing and analysis, but this requires more investment than depth colour coding of flat-mounted confocal images (Milde et al. 2013).

### Vascular Tuft Formation

The OIR model is a commonly used to study retinopathies. The three-dimensional nature of vascular tufts makes them ideal for LFSM and though this is a widely studied mouse model, the improved three-dimensional imaging allowed us to identify several new features of the important pathological vessels it generates. Our observations of small, medium and large tuft classes with distinct properties and the observation of more complex knotted, swirling and looping morphologies than previously reported, suggest a new mechanistic explanation is required to understand how and why vessels twist and turn on themselves and why it appears that medium tufts reach a critical size then stop twisting and instead coalesce into larger more stable structures, akin to the development of blood islands in retinal development (Goldie et al., 2008).

Nuclei with unusual shapes have previously been identified in abnormally growing tissues, such as cancer (Hida et al. 2004; Kondoh et al. 2013;, Versaevel et al. 2012), and to reflect mitotic instability (Gisselsson et al. 2001). It is remarkable that we observed the dramatically curved shape of EC nuclei in tufts. Although it remains unclear whether their unusual shape has consequences for EC function in the tuft, it is tempting to speculate that it would have some bearing on, or is at least be an indicator of abnormal cell behavior. Overall, the ability to rotate the tufts in 3D and view from the side, not just the top, gave a much clearer view of their structure potentiating a detailed analysis of their complex knotted structure in the future. LSFM therefore could greatly improve our understanding of these abnormal vascular formations, already opening up avenues for future studies.

Taken together, we propose that tuft formation proceeds as follows 1) small tufts are sprouts from the existing vasculature oriented upwards to the vitreous (e.g. as shown in Fig, 5a top panel) initiating the formation of a new tuft. However, further dynamic study is required to establish whether they are medium tufts that instead are regressing; 2) medium tufts originate from small tufts that continue to sprout, loop and potentially fuse with other upward sprouts (to generate the observed increase in connections to the plexus below), actively swirling and knotting around themselves. Further studies are needed to identify if other cell type/extracellular tissue acts as a scaffold to create the space/holes within them; 3) large tufts form when medium tufts merge with each other whereupon the knotting/swirling process declines (indicated by the plateau in curved nuclei in the large tufts and the sudden jump in tuft volume and connections to the plexus) possibly indicated final stabilization or even a switch to the natural normalization of tufts in this model.

### Reproducible Live imaging of angiogenesis in the retina

Current retinal studies must infer dynamics from static images by hypothesizing what might have happened in real-time to generate the retina’s phenotype. For example, CollagenIV-positive and IsolectinB4-negative vessels are considered to be empty membrane sleeves where the vasculature has regressed. It is therefore important to establish reproducible live-imaging methods. It will be interesting to investigate in future live-imaging studies how pervasive the kiss and run behaviors are across the plexus and under different conditions, in order to fully elucidate their functional role. We furthermore demonstrated the potential to quantify diverse subcellular level movements in the cells as proof of concept. The LSFM live imaging protocol is sturdy as indicated from the testing in three different laboratories in three different countries (US, Sweden and Portugal) with different scientists performing the dissections and imaging, on different instruments. As such we can confirm that though challenging, the live imaging protocol has been optimized and is reproducible in different hands.

## Materials and Methods

### Mice

mT/mG mice were crossed with cdh5 (PAC) CreERT2 mice (Wang et al. 2010). For live-imaging of retinal angiogenesis during development, mice were injected with 50 µg tamoxifen at postnatal day (P) 1, P2 and P3, and imaged at P4 (Wang et al. 2010). For live-imaging of oxygen-induced retinopathy (OIR) experiments, mice were injected with 100 µg tamoxifen at P13, P14 and P15. The retinal vasculature was imaged at P17. Recombination was confirmed by GFP expression in ECs. Mice used in experiments at Beth Israel Deaconess Medical Center were held in accordance with Beth Israel Deaconess Medical Center IACUC guidelines. Animal work performed at Uppsala University was approved by the Uppsala University board of animal experimentation. Transgenic mice were maintained at the Instituto de Medicina Molecular (iMM) under standard husbandry conditions and under national regulations.

### Antibodies

IsolectinB4 directly conjugated to Alexa488 and all corresponding secondary alexa conjugated antibodies were obtained from Invitrogen. Isolectin IB4 conjugated with an Alexa Fluor 568 dye was purchased from Thermo Fisher Scientific, MA. Anti-calretinin (ab702) and anti-ERG (ab2513) antibodies were obtained from Abcam. The antibody directed against Calbindin (AB1778) was acquired from Millipore. Anti-Glial Fibrillary Acidic Protein (GFAP) antibody was purchased from Dako (Z0334), anti-CollagenIV from AbD Serotec (2150-1470), biotinylated anti-neuron-specific b-III Tubulin from R&D Systems (Clone TuJ-1, BAM1195), and Cy3-conjugated anti-smooth muscle actin (SMA) antibody was obtained from Sigma Life Science (C6198). Draq5 was obtained from ThermoScientific. Anti-GOLPH4 (ab28049) from Abcam.

### Immunohistochemistry

Retinas were dissected as previously described (Del Toro et al. 2010). In brief, eyeballs were fixed for 18 min in 4% paraformaldehyde at room temperature. After dissection, retinas were blocked for 1 hour in blocking buffer (TNBT) or Claudio’s Blocking Buffer (CBB) for retinas with Golgi stained. CBB consists of 1% FBS (ThermoFisher Scientific), 3% BSA (Nzytech), 0.5% Triton X100 (Sigma), 0.01% Sodium deoxycholate (Sigma), 0,02% Sodium Azide (Sigma) in PBS pH = 7.4 for 2 hr in a rocking platform) for retinas stained for golgi. Thereafter, retinas were incubated overnight in primary antibody in blocking buffer. After extensive washing, retinas were incubated in the corresponding secondary antibody for 2 hours at room temperature. For confocal microscopy, retinas were mounted on glass slides, and for LSFM, retinas were mounted in 2% low-melting agarose. Agarose was melted at >65°C, and then maintained at 42°C before adding the tissue. To minimize curling of the retina, apply 1-2 drops of low melting agarose on retina and start uncurling the retina before the gel is solidified. It can then be transferred to the cylinder for imaging.

### PACT clearing of retinas

PACT clearing was performed as previously described (Treweek et al. 2015b). Retinas were dissected and fixed with 4% PFA at 4°C overnight. Samples were incubated overnight at 4°C in ice cold A4P0 (40% acrylamide, Photoinitiator in PBS). The following day, samples were degassed on ice by applying a vacuum to the tube for 30 min, followed by purging with N_2_ for 30 min. Thereafter, samples were incubated at 37°C for 3h to allow hydrogel polymerization. Excess gel was then removed from the samples, the samples washed in PBS, and incubated at 37°C for 6h in 8% SDS/PBS, pH 7.5. Samples were then washed in PBST for 1-2 days, changing wash buffer 4-5 times to remove all of the SDS. Immunostaining was then performed following the same protocol without PACT clearing. Thereafter, the tissue was cleared by at least 48h incubation in RIMS (40 g histodenz in 30 ml of sterile-filtered 0.02 M phosphate buffer, 0.01% sodium azide). Cleared retinas were mounted in 5% low-melting agarose/RIMS for LSFM imaging.

### Live-imaging

For live-imaging, mice were imaged at P4 or P5. The sample chamber was filled with DMEM without phenol red containing 50% FBS and P/S and heated to 37°C. Retinas were quickly dissected in prewarmed HBSS containing penicillin and streptomycin. After dissection, retinas were rapidly cut into quarters (mainly to minimize the datafile size created, the curved form was preserved) and immediately mounted in 1% low melting agarose in DMEM without phenol red containing 50% Fetal Bovine Serum (FBS) and 1x penicillin and streptomycin (P/S). To minimize curling of the retina, as with static imaging, apply 1-2 drops of low melting agarose on retina and start uncurling the retina before the gel is solidified. It can then be transferred to the cylinder for imaging.

### Equipment

All LSFM images were acquired with a Zeiss Z.1 light sheet microscope. The Zeiss objectives used for uncleared tissue and live imaging were Zeiss, RI=1.33, 5x/0.16, and 20x/1.0. For PACT cleared tissue, RI=1.45, 20x/1.0 (5.5mm working distance) was used. All raw data were handled on a high-end DELL workstation (Dual 8-core Xeon Processors, 196GB RAM, NVIDIA Titan Black GPU, Windows 7 64 bit) running ZEISS ZEN (Light sheet edition). Confocal images were taken with the LSM 880 Confocal Microscope.

### Image Analysis

#### Visualization of images in 3D

3D reconstructions of images up to 4 GB were obtained using Imaris software. Fiji was used for reconstruction of images larger than 4 GB. To quantify tuft volumes, Arivis Vision4D software was used.

#### Visualization of live-imaging

To visualize live-images, the maximum intensity projection of each timepoint was made in ZEN (Zeiss). The movies were corrected for drift correction in Fiji using the StackReg plugin and the background subtracted in Fiji using rolling ball background subtraction.

#### Deconvolution

In LSFM, when using a high NA (>∼0.6) objective, the optical section is determined by the depth of field of the objective and not the light sheet. However, in our system the light sheet is thicker than the objective’s depth of field and substantial out-of-focus light is captured relative to confocal. Additionally, thick tissue samples have an intrinsic milky appearance. This lack of clarity undermines sharp images and becomes progressively more of an impediment the deeper one tries to look into a tissue volume. This translucency is caused by heterogeneous light scattering (Richardson & Lichtman 2015). As the tissue used for imaging in LSFM is thick, fluorescent light originating from deep within the tissue is scattered during its travel through the tissue volume, back to the objective. This results in both in- and out-of-focus light arriving at an incorrect position on the camera causing objects to blur.

To deconvolve and reduce this light scatter computationally images were split into channels with their respective emission wavelength. Microscopic parameters (including pixel size, objective and excitation wavelength) were inserted into the settings of Huygens software for each channel followed by choosing the signal to noise ratio (SNR) for each image and run the deconvolution with the same settings. The resulted deconvolved images were inserted into Imaris for further analysis in 3D if needed. Huygens software was used for deconvolving all images.

#### Actin-rich bundles and filopodia tracking

Tracking was performed manually using ImageJ/Fiji. For filopodia tracking, each filopodia was tracked between each frame of imaging and different analysis was performed. For tracking the actin-rich bundles, the Manual Tracking plugin in Fiji was used to manually select the ROI (=region of interest) and follow the pathway of each trajectory. The trajectories and the pathway were overlaid. Each trajectory could be visualized using Montage function.

## Acknowledgements

We would like to thank Sven Terclavers (HCBI) for the excellent technical support and Joe Brock and the Illustration team at The Francis Crick Institute for aiding with 3D drawing of tuft knots. C.P and K.B. were supported by funding from Harvard Catalyst | The Harvard Clinical and Translational Science Center (National Center for Research Resources and the National Center for Advancing Translational Sciences, National Institutes of Health Award UL1 TR001102), the NEI (1R21EY027067-01), and BIDMC. K. B and P.A were supported by The Kjell and Märta Beijer Foundation. K.B. and L.C.W. were supported by a grant from the Knut and Alice Wallenberg foundation (KAW 2015.0030). L.V. was supported by a Victor A. McKusick fellowship from the Marfan Society. M.R. was supported by an EMBO fellowship (ALTF 2016-923). K.I.H. was supported by institutional training grant T32 HL07893 from the NHLBI of the NIH. L.V. was funded by the Victor A. McKusick Fellowship from the Marfan Foundation and BIDMC. D.F.C supported by EY025259, Lions Foundation, and NEI core grant P30 EY03790. K-S Cho: EY027067. C.A.F was supported by European Research Council starting grant (679368), the Fundação para a Ciência e a Tecnologia funding (grants: IF/00412/2012; PRECISE-LISBOA-01-0145-FEDER-016394; and a grant from the Fondation Leducq (17CVD03).

## Supplemental Figure Legends

**Supplemental Figure 1 related to Figure 1.**
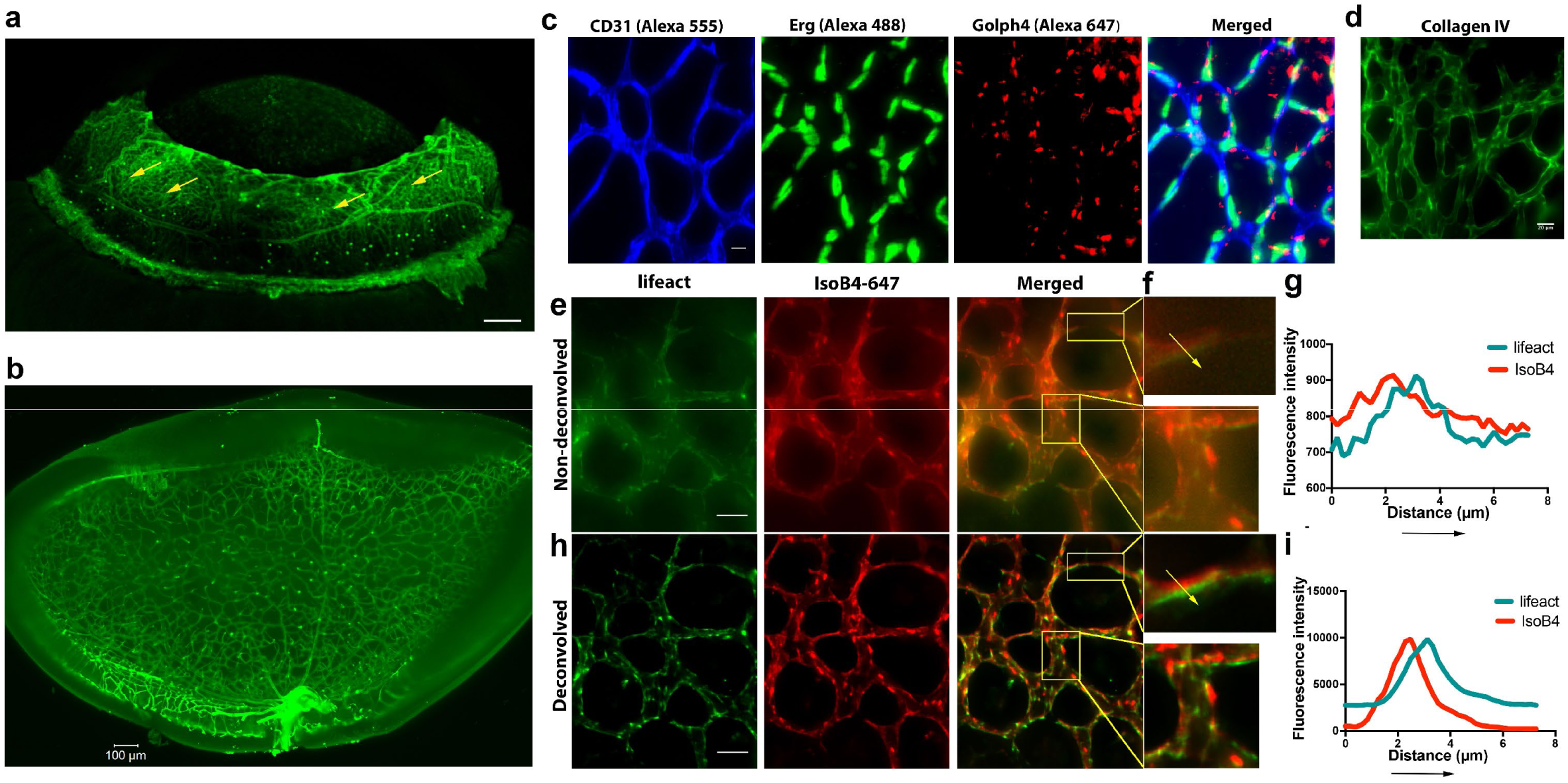
**a,** Maximum intensity projection of the iris microvasculature of two-week old mouse, LSFM. Vessels were visualized with IsoB4 (green). Scale bar, 200 µm; **b,** the full unprocessed retinal image used for the cropped figure in Fig. 1e. **c,** P12 retina stained using anti-CD31(Alexa 555), Anti-Erg (alexa488) and anti-Golgi (Golph4, Alexa 647) imaged using LSFM, Scale bar is 10 µm. **d,** P12 Retina stained for CollagenIV followed by LSFM imaging. Scale bar is 20 µm. **e,** lifeAct (green) mouse retinas stained for vessels (IsoB4 - red), merged images shows the overlap between these two channels together with the plot profile (**g**) of the selected area and the yellow arrow (**f**). **h,** the same channels were deconvolved using Huygens and **i,** the same area was plotted to compare the image resolution after deconvolution. Values were normalized to the maximum fluorescence intensity across both images at each pixel.

**Supplemental Figure 2 related to Figure 6.**
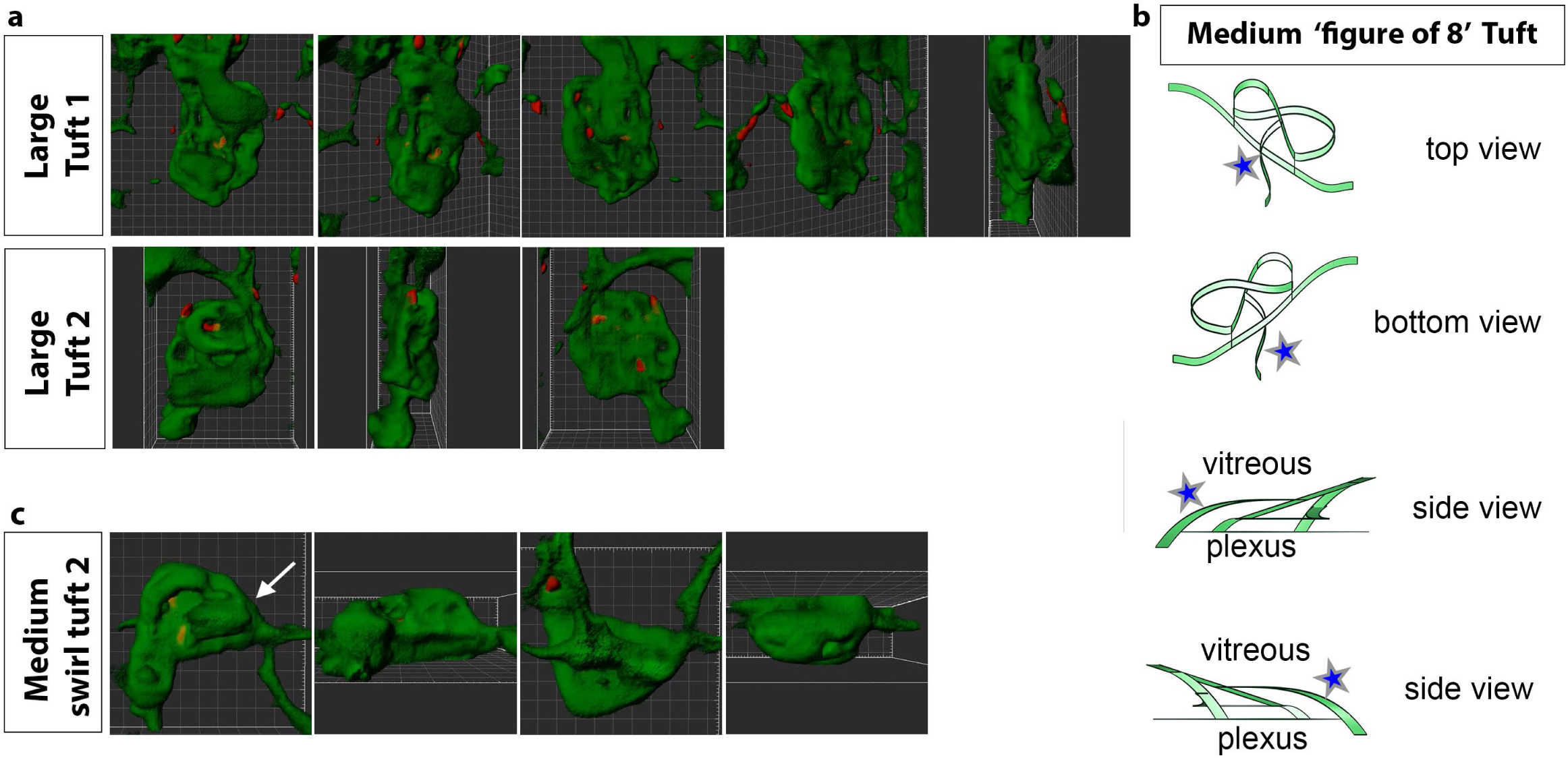
**a,** extended rotational views to see knotted structure from different angles of tuft 1 and 2 in Fig. 6a taken from movies S14, S15. **b,** 3D illustration (made with uMake software) while viewing the tuft with curved nuclei (from Fig. 6c and movie S13) from top to bottom with clipping planes to better understand how the knot topology changes through the tuft from the vitreous to the plexus side. Blue star indicates the unexpected vessel joining the top (vitreous facing) surface of the tuft to the plexus below. **c,** a second example of a medium tuft with a swirl structure and clear protrusive sprout-like morphology to the upper vitreous facing end point of the swirl.

## Movie Legends

**Movie S1.** 3D-rendered LSFM z-stack of retinal eye cups expressing yellow fluorescent protein, YFP (green) harvest from Thy1-YFP mice and stained with Isolectin IB4 (red).

**Movie S2.** 3D-rendered LSFM z-stack of all nuclei (Draq5, magenta) visualizes the inner nuclear layer and outer nuclear layer of the adult mouse retina. Vessels were visualized by IsolectinB4 staining (green). The retinal pigment epithelium emits green autoflourescence.

**Movie S3**. 3D-rendered LSFM z-stack of an adult mouse retina stained for Tuj1 (green) and horizontal cells (calbindin, red) visualizing the ganglion cells and horizontal cells, respectively.

**Movie S4.** 3D-rendered LSFM z-stack of an adult mouse retina stained for Smooth Muscle Actin (SMA, red) and CollagenIV staining (CollIV, green) visualizes the three vascular layers and smooth muscle cells in the mouse retina.

**Movie S5.** An hour time lapse LSFM movie showing long, slender filopodia, and rapid “kiss and run” fusion and disconnection of tip cells at the vascular front of mT/mG x cdh5 (PAC) CreERT2 mice. Frame rate: 1 image/45 sec.

**Movie S6.** A 9 hour time lapse LSFM movie showing a connection between two branches in the capillary plexus of mT/mG x cdh5 (PAC) CreERT2 mice. Frame rate: 1 image/20 min.

**Movie S7.** Representative tracking of a short-lived actin-rich bundle in life-Act-EGFP retina mice imaged with LSFM (7 min). The tracking was performed manually using Manual Tracking plugin in Fiji/ImageJ.

**Movie S8.** Representative tracking of a longer-lived actin-rich bundle in life-Act-EGFP retina mice imaged with LSFM (30 min). The tracking was performed manually using Manual Tracking plugin in Fiji/ImageJ.

**Movie S9.** Representative tracking of a long-lived actin-rich bundle track in life-Act-EGFP retina mice imaged with LSFM (40 min). The tracking was performed manually using Manual Tracking plugin in Fiji/ImageJ.

**Movie S10.** 3D-rendered LSFM z-stack of a ‘small tuft’ from an OIR mouse retina stained for blood vessels (IsolectinB4, green), and the nuclear marker ERG (magenta). The z-stack was rendered in Imaris.

**Movie S11.** 3D-rendered LSFM z-stack of a ‘medium tuft’ from an OIR mouse retina stained for blood vessels (IsolectinB4, green), and the nuclear marker ERG (magenta). The z-stack was rendered in Imaris.

**Movie S12.** 3D-rendered LSFM z-stack of a ‘large tuft’ from an OIR mouse retina stained for blood vessels (IsolectinB4, green), and the nuclear marker ERG (magenta). The z-stack was rendered in Imaris.

**Movie S13.** 3D-rendered z-stack showed a curved nuclei in a ‘medium tuft’ from an OIR mouse retina stained for blood vessels (IsolectinB4, green), and the nuclear marker ERG (magenta). The z-stack was rendered in Imaris.

**Movie S14.** 3D-rendered LSFM z-stack deconvolved in Huygens then reconstructed with surface rendering in Imaris of ‘Large tuft 1’ from an OIR mouse retina stained for blood vessels (IsolectinB4, green), and the nuclear marker ERG (red).

**Movie S15.** 3D-rendered LSFM z-stack deconvolved in Huygens then reconstructed with surface rendering in Imaris of ‘Large tuft 2’ from an OIR mouse retina stained for blood vessels (IsolectinB4, green), and the nuclear marker ERG (red).

**Movie S16.** 3D-rendered LSFM z-stack deconvolved in Huygens then reconstructed with surface rendering in Imaris of a ‘small tuft’ from an OIR mouse retina stained for blood vessels (IsolectinB4, green), and the nuclear marker ERG (red).

**Movie S17.** 3D-rendered LSFM z-stack deconvolved in Huygens then reconstructed with surface rendering in Imaris of ‘a Medium swirl tuft’ from an OIR mouse retina stained for blood vessels (IsolectinB4, green), and the nuclear marker ERG (red).

**Movie S18.** A 2.5 hour time lapse movie of a retinal tuft of a mouse in the OIR model using mT/mG x cdh5 (PAC) CreERT2 mice visualize short, rapidly extending and retracting filopodia as compared to control retinas. Frame rate: 1 image/minute.

**Movie S19.** A 2.5 hour time lapse movie of mouse retinal vasculature in the OIR model using mT/mG x cdh5 (PAC) CreERT2 mice reveal cell shuffling in real-time. Frame rate: 1 image/minute.

**Movie S20.** A 2.5 hour time lapse movie of mouse retinal vasculature in the OIR model using mT/mG x cdh5 (PAC) CreERT2 mice reveal abnormal vessel growth in real-time. Frame rate: 1 image/minute.

